# A Comparison of the Sexual Expression, Biomass, Cannabinoid Content, and Seed Production in XXX and XXY Triploid Cannabis

**DOI:** 10.1101/2024.09.25.612659

**Authors:** Nathan Paul, Ben Yanush, Adrian Monthony, Braydon Hall, Mehdi Babaei, Davoud Torkamaneh, AMP Jones

## Abstract

Cannabis is a multi-billion dollar industry reliant on unpollinated genetically female (XX) plants, as pollination reduces cannabinoid yields and flower quality. The mechanism behind the sexual expression of *C. sativa* is unclear. This study examines the sexual characteristics of homo- and hetero-gametic triploid cannabis plants (XX(X/Y)) to understand the role of the Y chromosome in cannabis sex determination. Heterogametic triploids were produced by crossing a male diploid *C. sativa* cv. Durban Poison (2n = 2x = 18 + XY) with a tetraploid female *C. sativa* cv. Higher Education (2n = 4x = 36 + XXXX) confirmed by flow cytometry and PCR. Measurements include fresh and dry plant weight, dry flower weight, seed counts, seed weights, and cannabinoids quantified using liquid chromatography. Results showed 44.4% of plants were XXY (*n* = 8), with 75% presenting a monoecious phenotype (*n* = 6). Of these, 50% produced predominately male flowers (*n* = 3), while the rest produced predominately female flowers. Plants with predominately female flowers had higher yields and cannabinoids compared to predominantly male plants. XXX genotypes outperformed XXY in several metrics. This study enhances our understanding of *C. sativa* sex determination and offers insight into breeding and cultivation strategies utilizing triploid cannabis.

## Introduction

*Cannabis sativa* L. (cannabis) is generally a short-day, diploid (2n = 20), annual, and herbaceous plant belonging to the Cannabaceae family (Raman *et al*., 2017; Small, 2017, 2022). Cannabis is primarily cultivated for its psychoactive secondary metabolites Δ9-Tetrahydrocannabinol (THC) and cannabidiol (CBD), fiber products, and seeds (Small, 2017). *C. sativa* can be monecious or dioecious, depending on the population (Moliterni *et al*., 2004; Ghosh *et al*., 2023). Many cultivars developed for fiber and seed production exhibit monoecious characteristics (Baldini *et al*., 2018), while genotypes used for recreational and medical purposes are predominantly dioecious with heteromorphic XX/XY sex chromosomes (Monthony *et al*., 2024). For the latter applications, producers usually cultivate exclusively female populations because they produce higher levels of secondary metabolites and to avoid unwanted pollination (Small, 2017; Punja & Holmes, 2020). The fertilization of females causes a reduction in secondary metabolites, as the plant devotes more resources to seed production (Small, 2017). This is of particular concern even when growing exclusively female populations as dioecious XX plants can produce male flowers, which poses a threat to the commercial production of medicinal cannabis (Punja and Holmes, 2020; Jones and Monthony, 2022).

Polyploidy refers to the genetic condition where more than two complete sets of chromosomes inhabit the same cell nucleus (Acquaah, 2012; Sattler, Carvalho and Clarindo, 2016). Polyploids that contain multiple sets of chromosomes from the same species are termed autopolyploids (Acquaah, 2012). On the contrary, polyploids that contain multiple sets of chromosomes from different species are termed allopolyploids (Acquaah, 2012). A significant effect of autopolyploidy is an increase in cell size (Acquaah, 2012). As a result, autopolyploid species typically display larger flowers, fruits, shoots and roots as well as broader, thicker leaves relative to their diploid counterparts (Acquaah, 2012; Sattler, Carvalho and Clarindo, 2016). In addition to enhancing organ size, polyploidy has been shown to contribute to increased levels of secondary metabolites in some plants, such as nicotine in tobacco (*Nicotiana tabacum* L.) (Acquaah, 2012). Another result of increased ploidy levels is a reduction in fertility, especially in odd ploidy levels (Köhler, Mittelsten Scheid and Erilova, 2010; Acquaah, 2012). This is primarily due to abnormal pollen formation and unbalanced meiosis and gametes causing sterility and/or premature seed abortion (Köhler *et al*., 2010; Acquaah, 2012). Progeny that do survive commonly contain unbalanced sets of chromosomes and are aneuploids (Köhler *et al*., 2010; Acquaah, 2012). Many species extensively used in commercial production are natural polyploids including banana (*Musa* sp.) cultivars (triploid) and alfalfa (*Medicago sativa* L.) (tetraploid) (Small Ernest & Marcel Jomphe, 1989; Acquaah, 2012; Sattler *et al*., 2016).

While polyploidy occurs naturally in many species, including cannabis, it can also be artificially induced (Philbrook *et al*., 2023). Due to the many desirable characteristics of polyploids, many plant breeders began inducing polyploidy on other diploid varieties through applications of exogenous mitotic inhibitors such as colchicine (Acquaah, 2012; Sattler *et al*., 2016). When applied to cells, colchicine disrupts spindle fibers in mitosis preventing the migration of recently duplicated chromosomes to opposite poles ultimately leading to a cell with double the normal sets of chromosomes (Acquaah, 2012; Sattler, Carvalho and Clarindo, 2016). Many species have been artificially induced into polyploids including seedless triploid watermelon (*Citrullus vulgaris* Schard.), and tetraploid snapdragons with larger flowers (*Antirrhinum majus* L. ‘Tetra Giant’; James Crow *et al*., 1994; Tolety & Sane, 2011).

While most cannabis is typically diploid, natural triploid and tetraploid varieties have been reported (Balant *et al*., 2022; Philbrook *et al*., 2023). Several groups have successfully induced polyploidization in both drug-type and hemp *C. sativa* (Bagheri and Mansouri, 2015; Parsons *et al*., 2019; Kurtz, Brand and Lubell-Brand, 2020, 2024; Crawford *et al*., 2021; Fernandes *et al*., 2023; McLeod *et al*., 2023; Tang *et al*., 2023; Suchoff *et al*., 2024). Polyploid cannabis plants are often larger and display an increased biomass relative to their diploid counterparts (Mansouri and Bagheri, 2017; Crawford *et al*., 2021; Fernandes *et al*., 2023; Suchoff *et al*., 2024). Polyploid cannabis has be shown to exhibit larger leaves, stems and larger but fewer stomata (Mansouri and Bagheri, 2017; Parsons *et al*., 2019; Kurtz, Brand and Lubell-Brand, 2020, 2024; Suchoff *et al*., 2024). The impact of polyploidy on cannabis secondary is highly variable (Bagheri and Mansouri, 2015; Mansouri and Bagheri, 2017; Parsons *et al*., 2019; Fernandes *et al*., 2023; Tang *et al*., 2023; Kurtz, Brand and Lubell-Brand, 2024). Some studies have shown increasing ploidy has no effect on THC production in cannabis (Bagheri and Mansouri, 2015; Parsons *et al*., 2019; Tang *et al*., 2023; Suchoff *et al*., 2024). Some studies have reported an increase in CBD as ploidy levels increased (Bagheri and Mansouri, 2015; Mansouri and Bagheri, 2017; Parsons *et al*., 2019; Crawford *et al*., 2021). However, others have indicated no significant differences in CBD production between ploidy levels (Tang *et al*., 2023; Kurtz, Brand and Lubell-Brand, 2024). Terpene concentrations have been found to highly variable in polyploid cannabis; terpene profile changes appear to be the most ubiquitous effect (Parsons *et al*., 2019; Tang *et al*., 2023; Kurtz, Brand and Lubell-Brand, 2024). Fernandes *et al*. (2023) showed that the response to increased polyploidy on secondary metabolite production in cannabis was highly cultivar dependent. Increased ploidy has been reported to result in fertility loss and seedlessness in cannabis (Crawford *et al*., 2021; Kurtz, Brand and Lubell-Brand, 2024; Suchoff *et al*., 2024). It has been shown that triploidy can reduce fertility between 77% to 100% relative to their diploid counterpart depending on cultivar (Crawford *et al*., 2021; Kurtz, Brand and Lubell-Brand, 2024; Suchoff *et al*., 2024). Crucially, studies have been limited to feminized or homogametic polyploid cannabis.

Most cannabis plants are photoperiodic, and flowering is induced under long nights with more than 12 hours of darkness, although the critical photoperiod varies among genotypes and day-neutral genotypes have been identified and developed (Moher, Jones and Zheng, 2021).

Drug-type cannabis is mostly dioecious, with distinct homogametic (XX) female and heterogametic (XY) male plants (Moliterni *et al*., 2004; Raman *et al*., 2017). Male inflorescences are arranged in clustered cymose panicles each containing five sepals, five stamens, and a pedicel (Raman *et al*., 2017). Female flowers comprise an ovary, style, and two stigmas enveloped in a perigonal bract, often covered in papilla cells and glandular trichomes at maturity (Spitzer-Rimon *et al*., 2019). These structures are arranged in condensed branchlets with a repeating phytomer structure, which collectively forms the *C. sativa* inflorescences often referred to as “buds” or “colas” (Shi, Schilling and Melzer, 2024; Spitzer-Rimon *et al*., 2019). Despite its dioecy, cannabis genotypes have been known to display a variable degree of hermaphrodism (Punja and Holmes, 2020). This divergence of floral sex from the chromosomal sex, or sexual plasticity, has been harnessed as the basis of the production of feminized seeds (XX seeds) and has been widely used in cannabis breeding programs (Lubell & Brand, 2018; Flajšman *et al*., 2021; Monthony *et al*., 2024). It has been noted that this can occur in response to stress, but for breeding purposes it is often induced through the application of ethylene inhibitors, or to a lesser extent gibberellic acid (Galoch, 1978; Mohan Ram & Sett, 1982; Monthony *et al*., 2024)

There are two primary mechanisms by which sex is determined in dioecious plants, an active Y system and an X-to-autosome ratio (Dellaporta and Calderon-Urrea, 1993). In an active Y system, sex is determined by the presence or absence of a Y chromosome where individuals possessing a Y chromosome are anatomically male and individuals absent of a Y chromosome are anatomically female (Dellaporta and Calderon-Urrea, 1993). The Y chromosome contains several sex determining regions (SDRs) that contribute to masculinization and suppression of female characteristics (Dellaporta and Calderon-Urrea, 1993). In this system, polyploid individuals with a single Y chromosome can overcome up to three X chromosomes and present as male (XXXY) (Dellaporta and Calderon-Urrea, 1993). The X-to-autosome ratio mechanism of sex determination relies on the ratio between the number of X-chromosomes to the total number of sets of autosomes (Shephard *et al*., 2000). In this system, SDRs can be found on X and Y chromosomes; interactions of these regions and autosomes ultimately determine sexual development (Dellaporta & Calderon-Urrea, 1993; Shephard *et al*., 2000). Polyploid species with a ratio of 1:1 (X:A) would be female, 0.5:1 male, and 0.5-1.0:1 hermaphroditic (Dellaporta and Calderon-Urrea, 1993). This method of sex determination is found in the closest relative to *C. sativa*, *Humulus Lupus* L. (hops) (Shephard *et al*., 2000). SDRs have been mapped to the X chromosome in hops (Clare *et al*., 2024); however, these markers may not be transferrable to *C. sativa* as it lacks the primer sequence present in commercial hop varieties (Clare *et al*., 2024). In addition, the marker failed to predict the sex of a wild relative of hops (Clare *et al*., 2024). Heterogametic triploid hops (XXY) have been shown to produce monoecious types when the X:A ratio is 0.66:1 (Haunold, 1971; Shephard *et al*., 2000). A study of the sexual expression of 575 triploid hops plants revealed that 70.6% produced exclusively female floral organs and 1.9% produced only male floral organs (Haunold, 1971). In addition, 18.1% of the triploids were predominately male monoecious types (about 5% female flowers), 6.3% of the triploids were monoecious with about even amounts of male and female flowers, and 2.1% were predominantly female monoecious types (about 95% female flowers) (Haunold, 1971).

Contrary to dioecious plants, the sex of monoecious species is typically controlled by a few sex-determining genes (Boualem *et al*., 2015). In *Zea Mays* L., the ts1 and ts2 genes regulate stamen and pistil production (Dellaporta and Calderon-Urrea, 1993). In the Cucurbitaceae family, a model system for the study of monoecious sex, floral sex in melon (*Cucumis melo*) is determined by a few alleles located at specific loci (Martin *et al*., 2009). Expression of *CmACS-7*, encoding for an ethylene biosynthesis enzyme ACS, suppresses the production of stamen in female flowers (Martin *et al*., 2009). Another ACS gene, *CmACS11* has also been demonstrated to be highly expressed in female flowers and hermaphrodite flowers but absent in male floral tissues (Boualem *et al*., 2015). *CmACS11* also acts as a repressor of melon *CmWIP1*, which when the latter is expressed alone leads to unisexual male flowers (Martin *et al*., 2009; Boualem *et al*., 2015). Sex determination of many cucurbits depends highly on the relative expression and repression of key ethylene-related genes (Martin *et al*., 2009; Boualem *et al*., 2015; Martínez & Jamilena, 2021). As such, the use of exogenous ethylene and its inhibitors has been used to control and optimize the ratio of female to male flowers to improve yield and to control production and labour constraints (Martínez and Jamilena, 2021; Oda *et al*., 2022).

There are few studies discussing the primary mechanism of sex determination in *C. sativa*. It has been proposed that sex is controlled through an X-to-autosome ratio where the Y chromosome contributes only a few key genes regulating sex expression (Grant *et al*., 1994; Adal *et al*., 2021). It has been reported that 35% of all sex-linked genes were mapped to autosomes whereas the remaining 65% were mapped to the sex chromosomes (Prentout *et al*., 2020). In contrast, a study exploring the role of ethylene-related genes in cannabis sexual plasticity found that 26.2% of Cannabis Ethylene Related Genes (CsERGs) were mapped to the sex chromosomes and the remainder located on the autosomes (Monthony *et al*., 2024). The study also only identified two genes located on the X chromosome, CsACO5 and CsMTN, with expression patterns associated with the sexual plasticity of cannabis (Monthony *et al*., 2024). The authors suggest that the sexual plasticity of *C. sativa* is controlled, in part, by the differential expression of CsERGs on both the sex chromosomes and autosomes (Monthony *et al*., 2024).

The gaseous hormone ethylene plays a crucial role in multiple signaling and response pathways in many plants and often impacts the sexual identity of flowers (Alonso and Stepanova, 2004). Ethylene receptors within plants such as ETHYLENE INSENSITIVE4 (EIN4), ETHYLENE RESPONSE SENSOR 1 (ERS1), ERS2, ETHYLENE RESISTANCE 1 (ETR1), and ETR2 require a copper co-factor for ethylene to bind (Alonso and Stepanova, 2004; Dubois, Van den Broeck and Inzé, 2018; Mckay, 2021). When ethylene fails to bind to these receptors, they interact with Raf-like kinase CONSTITUTIVE TRIPLE RESPONSE (CTR1) which in turn represses the positive regulator of ethylene ETHYLENE INSENSITIVE2 (EIN2) ultimately inhibiting ethylene production (Alonso and Stepanova, 2004; Mckay, 2021; Owen, Suchoff and Chen, 2023). It has been demonstrated that the application of exogenous silver-based ethylene blockers such as a mixture of silver thiosulfate and sodium thiosulfate (STS) triggers female cannabis plants to produce male inflorescences (Lubell & Brand, 2018; Owen *et al*., 2023). STS outcompetes copper ions at the binding sites, allowing CTR1 to remain active, thereby reducing ethylene production (Mckay, 2021). The ability to reverse sex by chemical and hormonal treatment suggests that the sexual characteristics of floral organs produced by floral primordia in *C. sativa* are transient and partially, continuously regulated (Dellaporta and Calderon-Urrea, 1993; Monthony *et al*., 2021).

This study aims to build upon previous knowledge of triploid *C. sativa* by examining the sexual expression, biomass, cannabinoid content and seed production of homo- and hetero-gametic triploid cannabis plants (2n = 3x = 27 + XX(X/Y)). If all individuals with a Y chromosome produce 100% male inflorescences and all individuals absent of a Y chromosome produce 100% female inflorescences, the mechanism of sex determination would likely be an active Y system. If it deviates from this and any individuals produce hermaphroditic flowers, it may point towards an X-to-autosome ratio system where it produces monoecious types at a ratio of 0.66-1:1. However, it may not follow either system like other species that rely on sex-determining loci as previously suggested by recent studies on the role of CsERGs in determining the phenotypic sex. Notably, this research represents one of the first investigation of XXX and XXY triploid drug-type cannabis plants. Investigating these characteristics is crucial for further advancing our understanding of the mechanisms behind sex determination and plasticity of *C. sativa*, as well improving breeding programs and cultivation practices. Ultimately, this study will further explore the potential of triploid cannabis utilization in commercial and medicinal applications.

## Materials and Methods

### Plant Materials

To obtain heterogametic triploid cannabis plants (2n = 3x = 27 + XXY), a male diploid *C. sativa* cv. Durban Poison (2n = 2x = 18 + XY; Dutch Passion, Netherlands) was crossed with a tetraploid female *C. sativa* cv. Higher Education (2n = 4x = 36 + XXXX; Remix Genetics, Canada). Sixty-two seeds were collected and germinated. After emergence, 18 plants were randomly selected and placed in a controlled growth chamber under an 18-hour photoperiod at 25°C. Plants were grown for 22 days before transitioning to a 12-hour photoperiod to induce flowering. The plants were flowered for 66 days before harvest. Following harvest, entire plants were hung upside down in the dark to dry in a controlled growth environment at a constant 25°C and 55% relative humidity.

### Ploidy Verification

To verify the ploidy level of all plants, flow cytometry was conducted on young leaf tissue collected before flowering. A 1cm^2^ piece of tissue was chopped via a razor blade into 1ml of nuclei extraction buffer. Samples were then filtered through a 20μm strainer where they were subsequently placed in a dark refrigerator (4°C). Samples were then analyzed on a BD FACSCalibur flow cytometer (BD Biosciences, San Jose, CA). Histograms were generated through the BD CellQuest Pro version 6 software (BD Biosciences, San Jose, CA). External standards of known diploid cannabis plants were used to verify the location of peaks. A soybean (*Glycine max* cv. “Polanka”) was used as an internal standard as it has a similar 2C DNA content (2.50 pg/2C) as *C. sativa* (1.97 pg/2C) (Doležel, Doleželová, and Novák, 1994; Parsons *et al*., 2019).

### Phenotype Classification

The ratio of female-to-male flowers was evaluated visually before harvest to determine phenotypes. The phenotype was based on the visual density of male and female floral organs. Plants were grouped into females with no male flowers, Predominantly Female Monoecious (PFM) with a majority (>50%) of female flowers, Predominately Male Monoecious (PMM), with a (<50%) minority of female flowers, and males with only male flowers. Images of whole plants and their branches were taken with an iPhone 12 Pro (Apple Inc., Cupertino, CA). Image backgrounds were removed using GIMP software (version 2.10.36).

### Floral Morphology Assessment

During flowering, clusters of inflorescences from each plant were collected and examined under a Zeiss Axio Zoom V16 macroscope (Zeiss, Oberkochen, Germany). Inflorescences were dissected under brightfield illumination to observe any abnormal development. Images were analyzed in ZEN 2.3 pro (Zeiss, Oberkochen, Germany).

### Pollen Germination Analysis

Pollen was collected from male inflorescences 2 weeks before harvest to assess pollen germination rates. A total of 10 male inflorescence clusters were collected and placed on pollen germination media (17% sucrose, 300 mg L^-1^ Ca(NO_3_)_2_, 100 mg L^-1^ H_3_BO_3_, and 0.7% agar, pH of 6.4) (Zottini *et al*, 1997). The samples were then placed under low-light conditions (∼50umol) for 18 hours to allow for pollen tube formation. The samples were then inspected to assess germination rates using a Zeiss Axio Zoom V16 macroscope. A 2mm x 2mm area was randomly selected to assess pollen germination percentages. The number of successfully germinated pollen grains was divided by the total number of grains in the image.

### Polymerase Chain Reaction (PCR) for Sex Determination

The prediction of cannabis individuals’ sex at the seedling stage was accomplished using a PCR-based assay as detailed by Borin *et al.,* (2021) and Törjék *et al.,* (2002) employing the following oligos: SCAR119_F: 5′-TCAAACAACAACAAACCG-3′ and SCAR119_R: 5′-GAGGCCGATAATTGACTG-3′. Subsequently, DNA fragment analysis was conducted, with a 1.5% agarose gel subjected to electrophoresis for 30 minutes at a voltage of 10V/cm in TAE buffer (0.4 M Tris acetate pH 8.3, 0.01 M EDTA). The gel was stained using SYBR safe DNA gel (Invitrogen, MA, USA) and subsequently visualized utilizing a gel imaging fluorescence system. Identification of male cannabis individuals was accomplished based on the presence of a 119 bp DNA fragment.

### Yield Measurements

The plants were harvested 5cm above the soil and weighed whole to gather fresh weights. After 3 weeks of drying, plants were weighed whole to gather dry weight and then processed into flowers. The flowers were trimmed following standard commercial practices and weighed for dry flower weight. The Harvest Index (HI) was calculated by dividing the dry flower weight by the whole plant’s dry weight. Any seeds produced by the plants were removed after the dry flower was weighed. Seeds were counted and weighed on a per-plant basis. Only whole seeds were included in the measurement.

### Cannabinoid Analysis

To determine the cannabinoid contents of the flowers, approximately 1 gram samples were randomly collected from each plant. Cannabis flower extraction was completed according to Mudge, Murch and Brown (2017) with modifications. Dried cannabis flowers were submerged in liquid nitrogen in a ceramic mortar and ground with a pestle until a fine powder was formed and the nitrogen was evaporated. 0.5g of material was sampled into a 50mL falcon tube, followed by 20mL of 99% ethanol. Samples were sonicated at room temperature for 15 minutes. Samples were centrifuged for 2 minutes at 350 RCF and supernatant decanted into a second falcon tube. The extraction was repeated with 20mL more of ethanol with sonication and centrifugation. Supernatants were pooled and filtered through a 0.22µm PTFE syringe filter, collecting 1mL in amber glass HPLC vials.

Extracts were analyzed using a Shimadzu Prominence-i LC-2030C Plus Liquid Chromatograph equipped with a UV detector. A Phenomenex Kinetex C18 column 100 x 2.1 mm, 2.6µm particle size, with a Phenomenex C18 guard column was used for separation. Mobile phases consisted of water (A), acetonitrile (B), and methanol (C), with 0.1% formic acid in each. The gradient elution method was initiated at 30%B and 35%C, then increased to 40%B and 30%C over 8 minutes, then to 100%B over the next 6 minutes where it was held for a further 3 minutes. Mobile phases were then returned to initial conditions for a 4 minute re-equilibration. The flow rate was set to 0.45ml/min with column temperature set to 37°C. The detection and quantitation wavelength was set to 230nm, with peaks matching retention times of certified reference standards (Certilliant, Austin, TX).

### Statistical Analysis

Data were analyzed utilizing two separate one-way ANOVA tests with genetic composition at the sex locus (XXY or XXX) and phenotypes (Male, PMM, PFM, and Female) as independent factors. The response variables, fresh weight, dry weight, dry flower weight, average Tetrahydrocannabinol acid (THCA) plus THC, number of seeds, average seed weight and harvest index, were separately tested. In addition, the data was fit to a linear regression model to analyze the relationship between increasing feminization and our response variables. Analysis post-hoc utilized a Tukey Honest Significant Differences (HSD) test to compare means of phenotypic groups. To compare means of heterogametic and homogametic plants post-hoc, a Welch’s two-sample t-test was used. Significance was declared at p <0.05 for all tests. Mean and standard errors were recorded for each treatment. Data and plots were analyzed and generated in R Studio. Phenotypic males were removed from all statistical analysis due to lack of replication available; their means are included in the means comparison table for reference (*n* = 2). Homogametic plants that displayed PFM phenotypes were also removed from the statistical analysis due to a lack of replication available (*n* = 2).

## Results

### Ploidy Verification

Flow cytometry revealed that 100% of plants were triploid (2n = 3x = 30) (n = 18).

### Phenotype Classification

Sex scoring was performed once 100 % of plants (*N = 18)* displayed floral organs after 12 days (Fig. 1). In total, 44.4% of individuals displayed exclusively female floral organs (*n* = 8) (Fig. 2*A*), 44.4% were monoecious with varying ratios of male, female and bisexual flowers (*n* = 8)(Figs. 2*B*-*F*), and 11.1% displayed exclusively male floral organs (*n* = 2 ) (Fig. 2*G*). The ratio of female to male flowers varied greatly within the individuals displaying both male and female floral organs. Of all plants, 16.7% displayed a PMM phenotype (*n* = 3) (Figs. 2*D*-*F*) and 33.3% displayed a PFM phenotype (*n* = 6) (Figs. 2*B* and 2*C*).

**Fig. 1.**
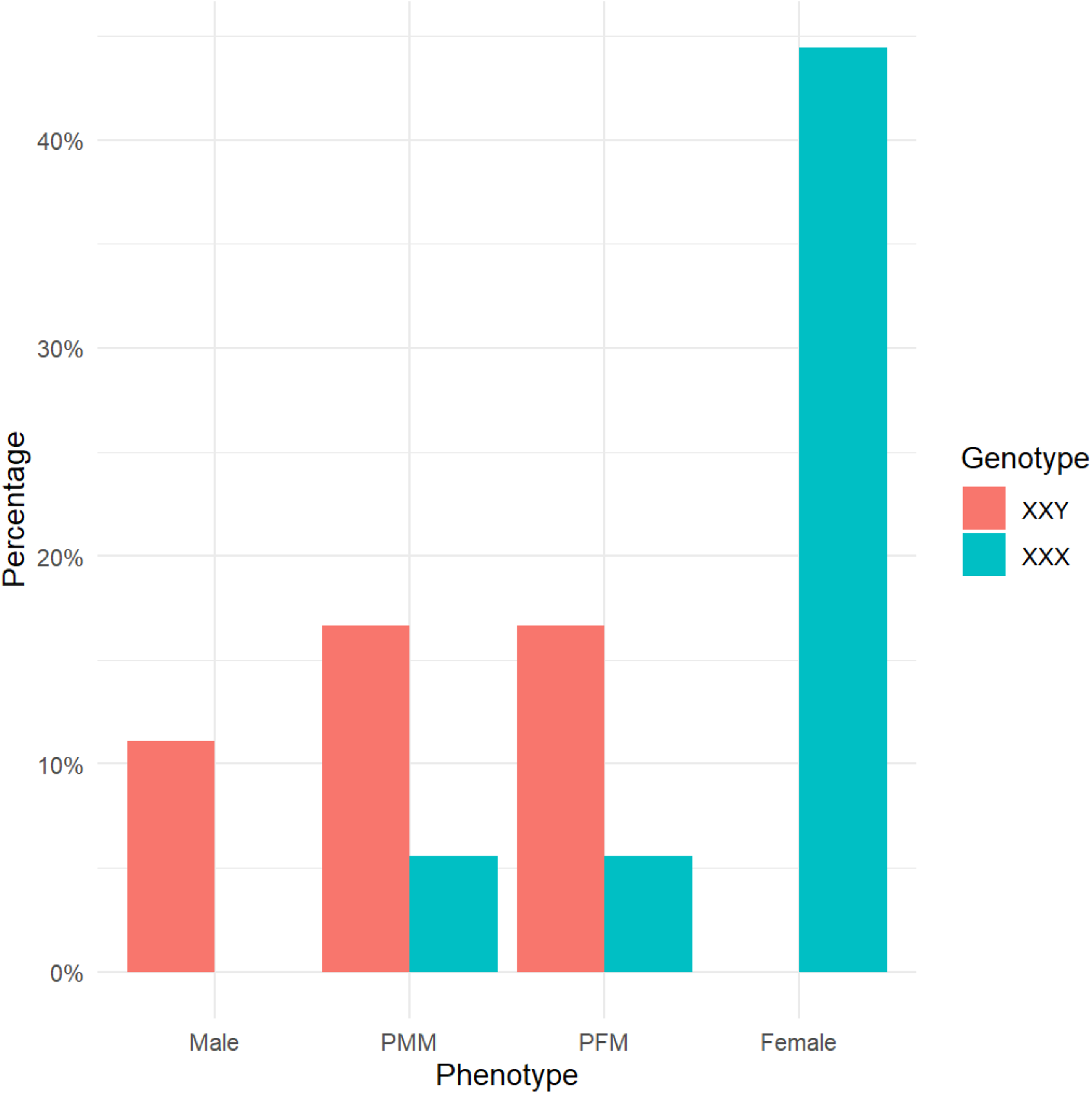
Phenotypic distribution of homo (*n* = 8) and heterogametic (*n* = 10) triploid *Cannabis sativa* L. plants. Predominantly Male Monoecious (PMM), Predominately Female Monoecious (PFM). Y-Axis represents the percentage of individuals of the total experimental population (combined XXY and XXX plants; *N* = 18).

**Fig 2.**
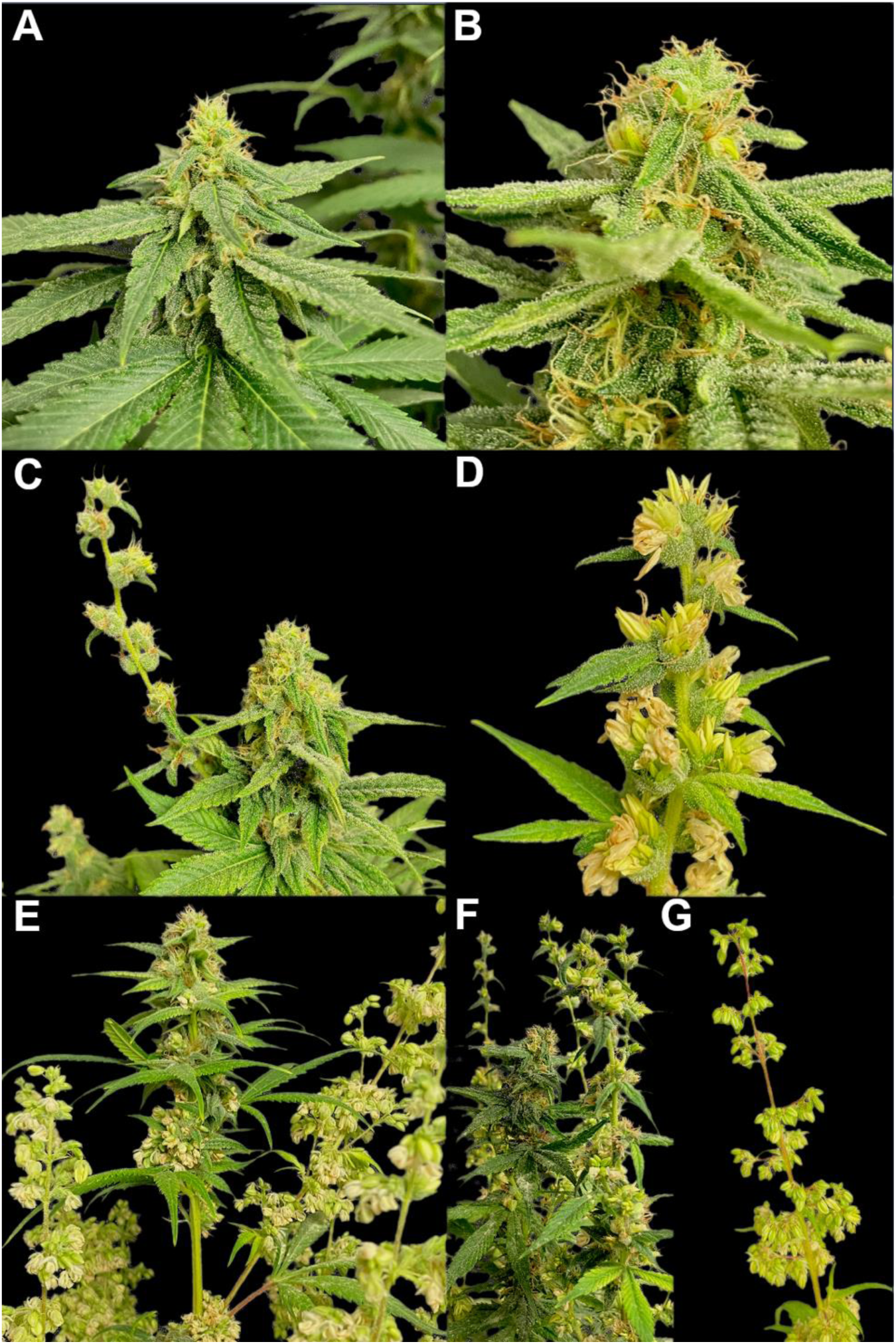
Representative photos of triploid inflorescences of *Cannabis sativa* L. 8 weeks after flower initiation. (A) Homogametic triploid (XXX) inflorescence displaying only female florets. (B-C) Examples of heterogametic triploid (XXY) inflorescence with a PFM phenotype. (D-F) Examples of heterogametic triploid (XXY) inflorescence with an PMM phenotype. (G) Heterogametic triploid (XXY) inflorescence displaying only male florets.

### Floral Morphology Analysis

The distribution and anatomy of flowers varied greatly among monoecious individuals (Figs. 2*B*-*F*). Some plants exhibited a separation of morphologically typical male and female flowers among branches (Figs. 3, 4*A* and 4*B*), while others showed clustering of these flowers next together (Figs. 4*C* and 4*D*). Many of the male flowers were atypical, with stamen emerging from female-like perigonal bracts (Figs. 4*A*, 4*B*, and 4*D*). The majority (66.7%) of all XXY monoecious plants (*n* = 6) produced at least one atypical bisexual flower with both stamen and pistils emerging from a perigonal bract (Fig. 5). Apical branches tended to be dominated by female floral organs, whereas lateral branches often produced primarily male flowers (Fig. 3). Notably, it was observed that apical portions of branches with exclusively male flowers tended to have more stamen emerging from female-like bracts (Fig. 3).

**Fig 3.**
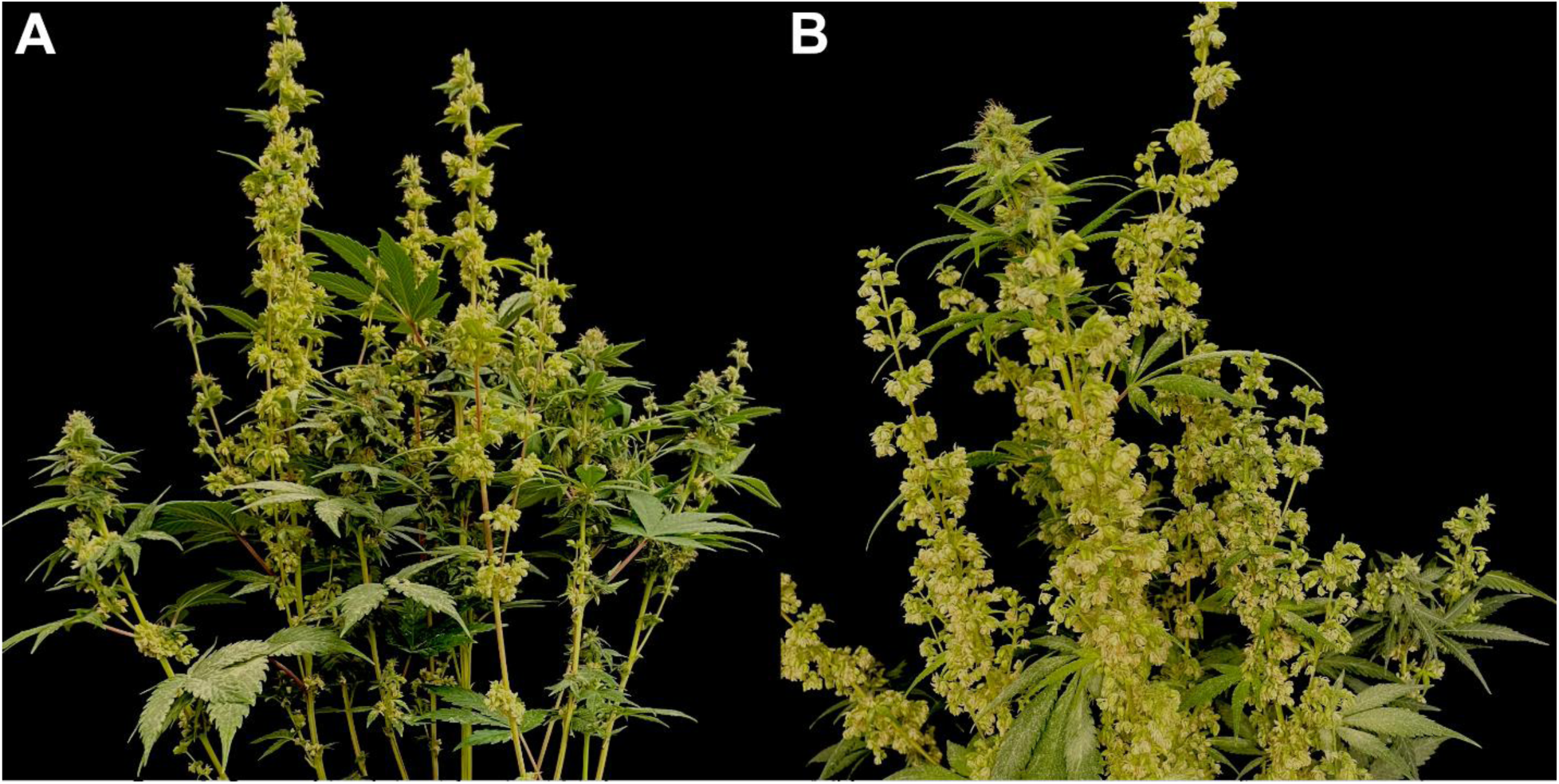
Representative photos of two heterogametic (XXY) triploid *Cannabis sativa* L. plants 8 weeks after flower initiation with separate male and female inflorescences. (A) Displays inflorescences with higher rates of female floral organs on axillary branches originating from the first node. (B) Shows an apical inflorescence with predominantly female floral organs. Axillary branches primarily bear male inflorescences, some of which protrude from bract structures, while apical branches mainly produce inflorescences with female characteristics. Apical male inflorescences tended to produce atypical male flowers emerging from female-like bracts.

**Fig 4.**
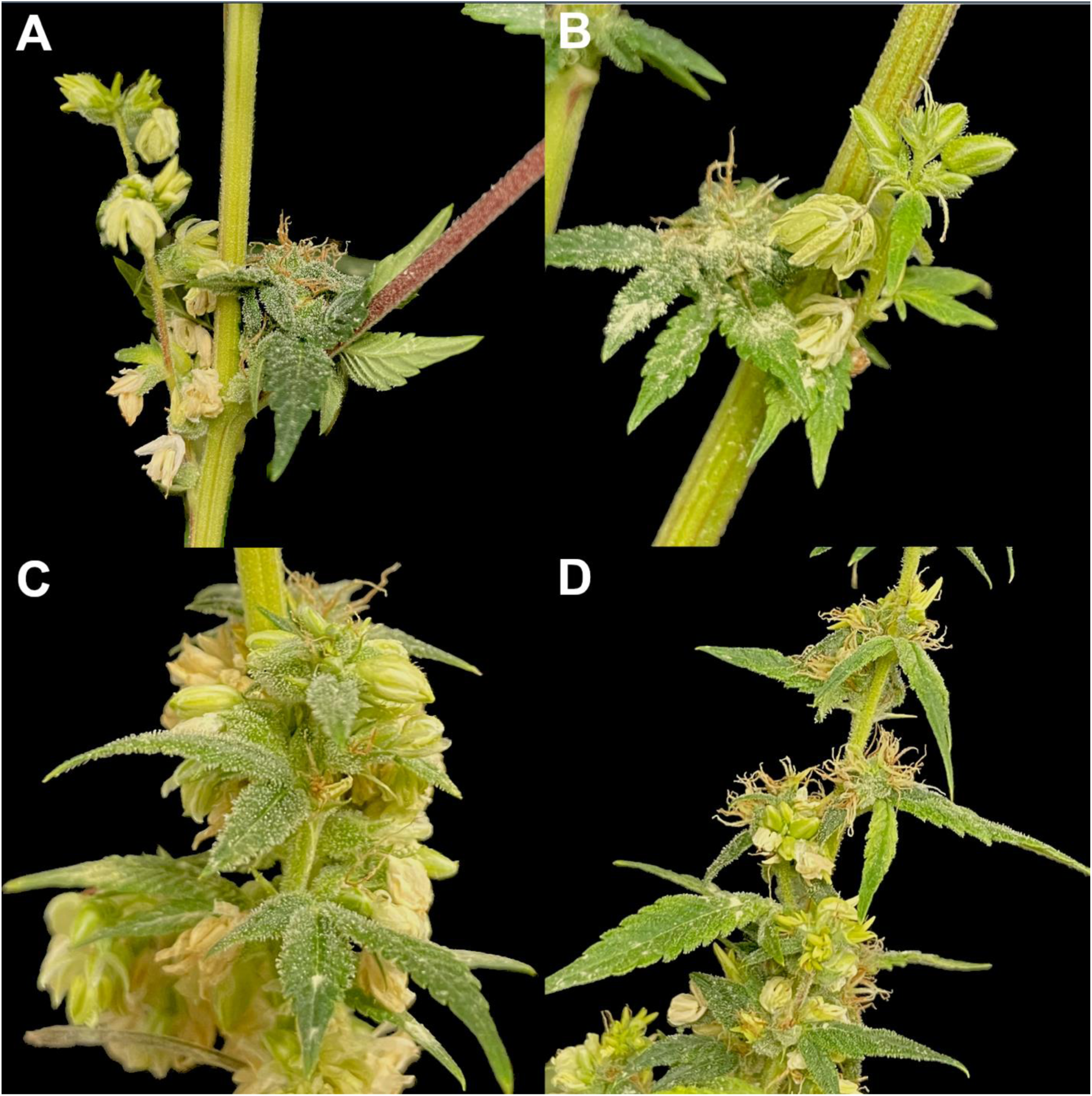
Representative photos of four heterogametic (XXY) triploid *Cannabis sativa* L. plants producing male and female inflorescences on the same branches 8 weeks after flower initiation. (A-B) Clusters of male inflorescences adjacent to female inflorescence and (C-D) inflorescences that have both male and female flowers adjacent to each other.

**Fig 5.**
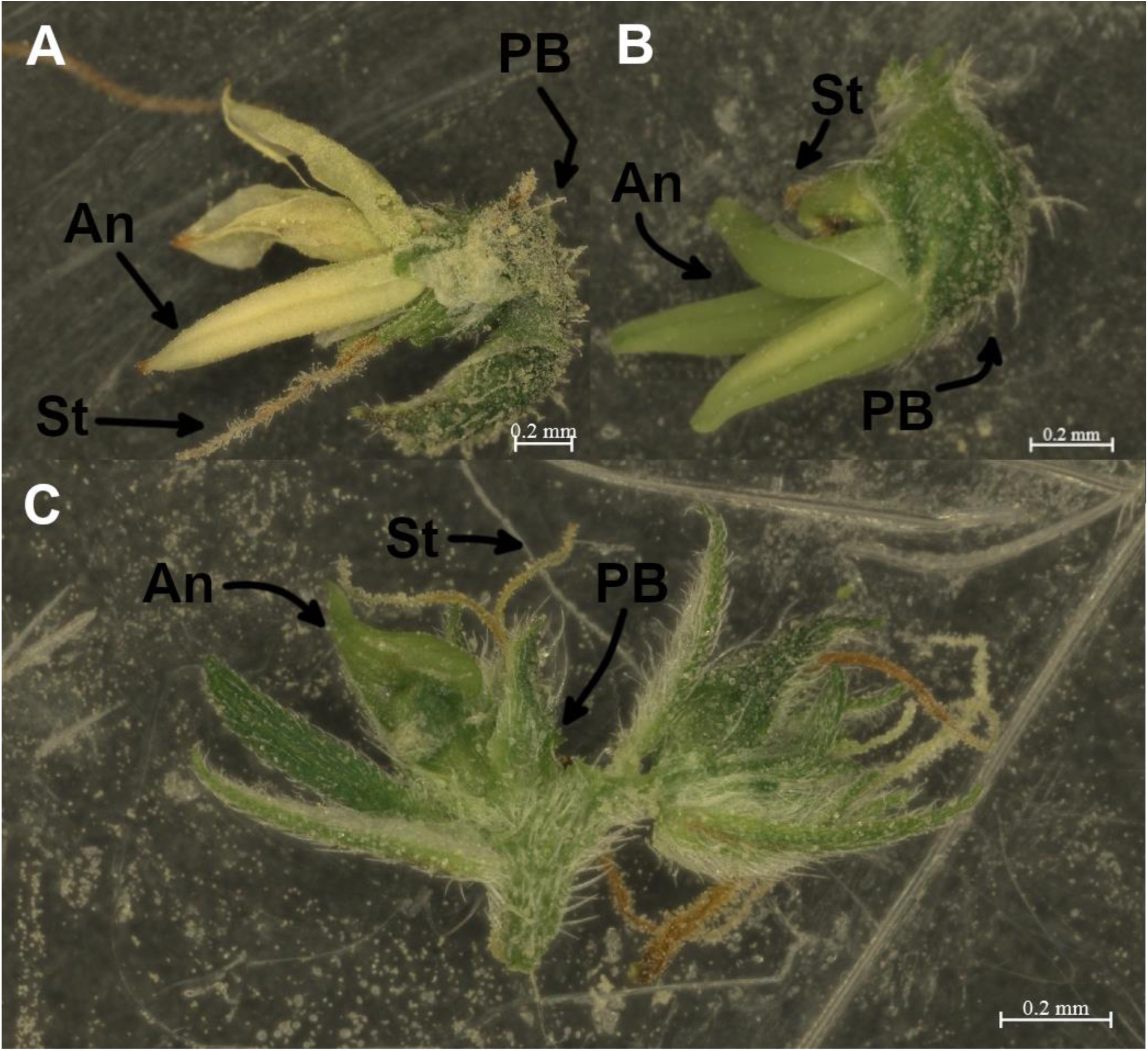
Stereomicroscopic images of bisexual *Cannabis sativa* flowers. (A-B) Single cannabis flowers displaying both two Stigma (St) and four Anthers (An) emerging from a single Perigonal Bract (PB). (C) Inflorescences displaying both male and female flowers.

### Pollen Germination Assay

Pollen germination rates were observed to be zero across all individuals that produced male floral organs and failed in 90% of individuals. In the one exception, one plant exhibiting both male and female floral organs had a pollen germination rate of <1%. Pollen production was highly variable among all pollen-producing individuals with some producing large amounts and others little to none.

### PCR Sex Determination Assay

Results from the PCR sex determination assay revealed that 44.4% of individuals carried a Y chromosome (*n* = 8). The remaining 66.6% of individuals were homogametic (*n* = 10). Of the XXY plants, 75% displayed a monoecious phenotype (n = 6). Of the monoecious XXY plants, 50% displayed a PFM phenotype (*n* = 3), and 50% displayed a PMM phenotype (*n* = 3). However, 20% of homogametic individuals produced monoecious phenotypes (*n* = 2).

### Yield Measurements

Fresh weight was significantly affected by phenotype (p = 0.008) and genotype (p = 0.014). The fresh weight of phenotypic females and predominantly female monoecious phenotypes was found to be significantly higher than predominant male monoecious phenotypes (p = 0.007, p = 0.033) (Table 1). The fresh weight of XXY genotypes was found to be significantly lower than XXX genotypes (p = 0.017) (Table 2). In contrast, neither phenotype nor genotype had a significant effect on dry plant weight at p <0.05 (p = 0.051, p = 0.079).

**Table 1.**
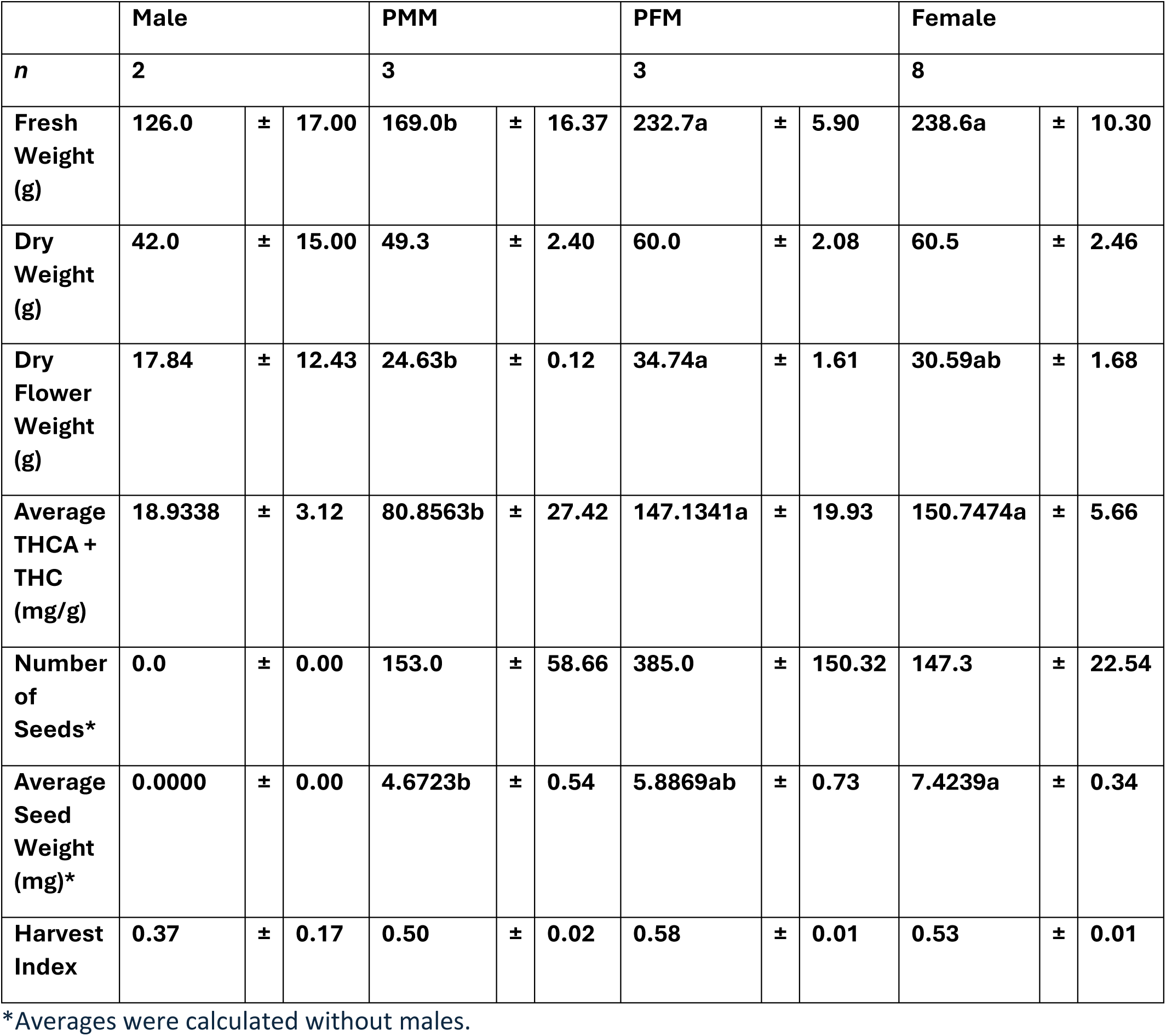
Means comparison and standard errors of four phenotypes (Male, PMM, PFM, and Female) of 16 triploid *Cannabis sativa* plants across growth metrics: fresh weight (g), dry weight (g), dry flower weight (g), average THCA+THC (mg/g), average seed weight (mg), and harvest index after 66 days of flowering. A mean comparison with a Tukey Honestly Significant Difference (HSD) test was generated post-ANOVA in R. Statistical significance was determined at p<0.05. Alternate lowercase letters indicate significant differences in means at p <0.05, as determined by the Tukey HSD Test.

|  | Male |  |  | PMM |  |  | PFM |  |  | Female |  |  |
| --- | --- | --- | --- | --- | --- | --- | --- | --- | --- | --- | --- | --- |
| *n* | 2 |  |  | 3 |  |  | 3 |  |  | 8 |  |  |
| Fresh Weight (g) | 126.0 | ± | 17.00 | 169.0b | ± | 16.37 | 232.7a | ± | 5.90 | 238.6a | ± | 10.30 |
| Dry Weight (g) | 42.0 | ± | 15.00 | 49.3 | ± | 2.40 | 60.0 | ± | 2.08 | 60.5 | ± | 2.46 |
| Dry Flower Weight (g) | 17.84 | ± | 12.43 | 24.63b | ± | 0.12 | 34.74a | ± | 1.61 | 30.59ab | ± | 1.68 |
| Average THCA + THC (mg/g) | 18.9338 | ± | 3.12 | 80.8563b | ± | 27.42 | 147.1341a | ± | 19.93 | 150.7474a | ± | 5.66 |
| Number of Seeds* | 0.0 | ± | 0.00 | 153.0 | ± | 58.66 | 385.0 | ± | 150.32 | 147.3 | ± | 22.54 |
| Average Seed Weight (mg)* | 0.0000 | ± | 0.00 | 4.6723b | ± | 0.54 | 5.8869ab | ± | 0.73 | 7.4239a | ± | 0.34 |
| Harvest Index | 0.37 | ± | 0.17 | 0.50 | ± | 0.02 | 0.58 | ± | 0.01 | 0.53 | ± | 0.01 |

\*Averages were calculated without males.

**Table 2.** Means comparison and standard errors of 8 heterogametic (XXY) and 8 homogametic (XXX) triploid Cannabis sativa plants across growth metrics: fresh weight (g), dry weight (g), dry flower weight (g), average THCA+THC (mg/g), average seed weight (mg), and harvest index after 66 days of flowering. Post-ANOVA, a Welch’s two-sample t-test was used to compare means. All analysis was conducted in R. Statistical significance was determined at p<0.05. Alternate lowercase letters indicate significant differences in means at p <0.05, as determined by the Tukey HSD Test.

|  | XXY |  |  | XXX |  |  |
| --- | --- | --- | --- | --- | --- | --- |
| *n* | 8 |  |  | 8 |  |  |
| **Fresh Weight (g)** | **182.1b** | ± | **17.00** | **238.6a** | ± | **16.37** |
| **Dry Weight (g)** | **51.5** | ± | **15.00** | **60.5** | ± | **2.40** |
| **Dry Flower Weight (g)** | **26.72** | ± | **12.43** | **30.59** | ± | **0.12** |
| **Average THCA + THC (mg/g)** | **90.2299b** | ± | **3.12** | **150.7474a** | ± | **27.42** |
| **Number of Seeds** | **201.8** | ± | **0.00** | **147.3** | ± | **58.66** |
| **Average Seed Weight (mg)\*** | **5.2796b** | ± | **0.00** | **7.4239a** | ± | **0.54** |
| **Harvest Index** | **0.50** | ± | **0.17** | **0.53** | ± | **0.02** |

\*Test was conducted on only seed-bearing XXY plants ( $n = 6$ ) and XXX plants ( $n = 8$ ).

Phenotype had a significant effect on dry flower weight while genotype did not (p = 0.029, p = 0.336). Predominant female monoecious were found to produce significantly higher dry flower yields relative to predominant male monoecious types (p = 0.024) (Table 1).

Both phenotype and genotype were found to have significant effects on average THC + THCA content (p = 0.010, p = 0.018). Predominant male monoecious phenotypes were found to have significantly lower average THC + THCA compared to phenotypic females and predominantly female monoecious phenotypes (p = 0.009, p = 0.036) (Table 1). When comparing genotypes, it was found that the XXX genotypes produced significantly higher levels of THC + THCA on average (p = 0.023) (Table 2). The highest average total THC and THCA (17.9%) was found in a predominantly female monoecious phenotype. The lowest average THC and THCA (1.6%) was found in a phenotypic male.

Neither phenotype nor genotype significantly affected seed production when not considering phenotypic males (p = 0.052, p = 0.155). The highest number of seeds produced was 593. The least amount produced that was not a phenotypic male was 6. Predominately female monecious plants were found to produce significantly more seeds than females (p = 0.049) (Table 1). However, the production of seeds was highly variable in the XXYs and monoecious plants, especially predominately female monoecious.

Both the phenotype and genotype were found to have a significant effect on average seed weight (p = 0.006, p = 0.003). Phenotypic females were found to have significantly heavier seeds on average when compared to the predominately male monoecious phenotypes (p = 0.006) (Table 1). In addition, the XXX genotypes had significantly heavier seeds on average relative to the XXY genotypes (p = 0.005). It was observed that many seeds produced were hollow, immature, and poorly developed (Fig. 6).

**Fig 6.**
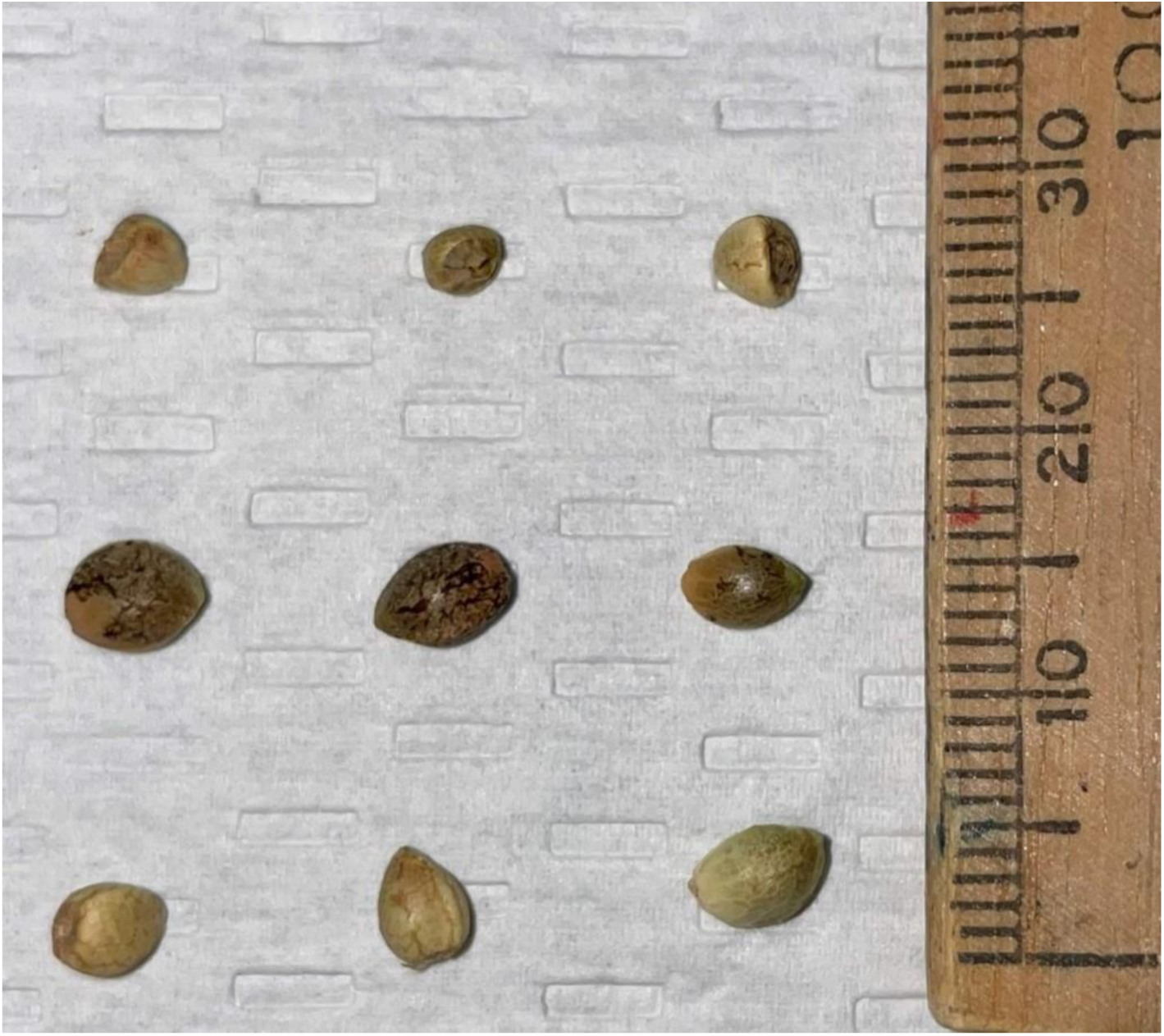
Seeds were collected from 14 homogametic (n = 8) and heterogametic (n = 6) triploid Cannabis sativa plants. Seed quality and variability varied considerably throughout. Some seeds were observed to be hallowed and malformed (Top row) while others were immature (Bottom row). However, it was observed that some seeds developed normally and did not have any malformities (Middle row). Seeds of all qualities were found in both homogametic and heterogametic plants.

Neither phenotype nor genotype were found to have a significant effect on HI (p = 0.075, p = 0.530).

## Discussion

To further the understanding of the mechanisms behind sex determination in *C. sativa*, this study examines the phenotypic expression of sex in hetero and homo gametic triploid cannabi*s.* Of the heterogametic (XXY) individuals, the majority were observed to produce both male and female reproductive organs at various ratios. Likewise, while Homogametic (XXX) triploids produced primarily female organs, low levels of male florets were also observed. Hermaphrodism is not a rare occurrence in cannabis; in a study of 1,000 female cannabis plants flowered for 6-7 weeks, 5-10% of the plants exhibited some degree of hermaphrodism by producing male florets (Punja and Holmes, 2020). While the rates observed here were slightly higher (20% of XXX plants), it is not known if this was a result of triploidy, the genetic background of the plants used, or the small sample size (*n* = 10). Few studies have been conducted on the natural hermaphrodism of male (XY) *C. sativa*. However, Moon *et al*. (2020) reported genotypic male (XY) plants treated with ethephon, a chemical agent that converts to ethylene gas, that closely resembles the XXY plants generated in this study. This sexual plasticity in genetically female plants and heterogametic (XXY) triploids, demonstrates that while the X and Y genes drive sexual expression, they interact with other factors to determine the ultimate final floral identity.

The effect of exogenous ethylene blockers and growth regulators such as gibberellic acid and auxin on sex expression is relatively well understood in *C. sativa* (Heslop-Harrison, 1956; Galoch, 1978; Mckay, 2021; Flajšman *et al*., 2021; Owen *et al*., 2023). These groups of compounds are known to induce the formation of male florets in genetically female plants, while the application of ethylene can induce female florets to develop on genetically male plants (Galoch, 1978; Moon *et al*., 2020). These observations demonstrate that in addition to the X/Y genes, various plant growth regulators are involved in determining the sexual identity of florets. Some of the monoecious plants produced in this study had a strong resemblance to diploid female *C. sativa* treated with ethylene blockers such as STS. For example, sub-optimal induction of male florets in genetically female diploid cannabis plants often results in bisexual florets, where stamens are produced within bracts that anatomically resemble female florets (Flajšman *et al*., 2021). While these stamens often do not shed pollen, they represent a partial sex reversal similar to what was observed in some XXY plants (Mejía Londoño, Barrera-Sánchez and Córdoba Gaona, 2023). The variability in the ratio of male to female flowers in heterogametic triploids indicates that sex is not determined merely by the presence of a Y chromosome in *Cannabis sativa*. Moreover, these results suggest sex is not strictly determined by an X to autosome ratio as previously reported by Grant *et al.,* (1994) as it involves other genetic and environmental factors. This is supported by previous research where a substantial amount of sex determining genes were mapped to autosomes and were not restricted to the sex chromosomes (Prentout *et al*., 2020; Adal *et al*., 2021; Monthony *et al*., 2024).

The interaction of X/Y genes with internal hormonal signalling also likely contributed to the heterogeneity of sexual expression in many of the heterogametic XXY triploids in this study. For example, many plant growth regulators are synthesized in specific regions and establish gradients within the plant to drive plant development and architecture (Bhalerao and Bennett, 2003; Petersson *et al*., 2009). One of the most well studied is auxin, which is produced in the apical meristem and leaves and moves toward the proximal end of the plant to form a concentration gradient (Bhalerao and Bennett, 2003). These gradients can drive plant development to help adapt to various situations (Bhalerao and Bennett, 2003). In the case of sex identity within heterogametic (XXY) cannabis plants, these gradients are likely responsible for the uneven distribution of male/female florets within individual plants. It was noted that apical portions of the inflorescences tended to consist of primarily female florets interspersed with some male florets, and it was quite common to different inflorescences on a single plant with different male to female floret ratios. Previous work has demonstrated that auxin promotes female flower development in cannabis, which could explain the increased prevalence of female florets in the apical regions observed in this study (Galoch, 1978). Another interesting observation was the variability in sexual expression in heterogametic (XXY) individuals. While some were phenotypically male (*n* = 2) the majority (*n* = 6) were monoecious to varying degrees (Fig. 1). Some plants displayed PFM phenotypes and were comprised of almost exclusively female flowers with some non-pistillate bisexual and male flowers (Fig. 2*B*) while the PMM phenotypes produced mostly non-pistillate bisexual flowers and male flowers and few female flowers (Fig. 2*D*). It was observed that within PMM and PFM plant groups, there was some variability in the amount of male, female and bisexual flowers and the distribution across the plant. It can be argued that this expression of sex is closer to what has been observed in monoecious hemp where individual plants vary in the ratio of male to female florets (Baldini *et al*., 2018). As such, it appears that while cannabis loosely follows the X to autosome sex determination model, it is further modulated through hormonal factors, which can in turn be impacted by environmental cues. The gradient of sexual expression that was observed, instead of discrete categories, suggests that there are multiple genes involved in this trait and there is significant diversity within this population.

In commercial production, all plants are female as male plants are destroyed (Punja and Holmes, 2020). Seed formation reduces female inflorescence quality; thus, producers must eliminate pollen-producing genotypes (Punja and Holmes, 2020). It has been reported that diploid hermaphroditic pollen germination rates were variable, ranging from 10%-30% (Punja and Holmes, 2020). The low pollen germination rates observed in this study are similar to what has been observed in triploids in other species (Zhang *et al*., 2016; Anamthawat-Jónsson *et al*., 2021; Subasinghe Arachchige *et al*., 2022). However, even extremely low germination rates do not mean that the plant is completely infertile. Several groups have demonstrated that triploid cannabis is not completely sterile and can produce seeds with both diploid and tetraploid pollen donors (Crawford *et al*., 2021; Kurtz, Brand and Lubell-Brand, 2024; Suchoff *et al*., 2024). Triploid species are typically rendered infertile due to the unbalanced gametes causing seed abortion shortly after fertilization (Köhler, Mittelsten Scheid and Erilova, 2010). However, some progeny may survive with an unbalanced set of chromosomes as aneuploids (Köhler, Mittelsten Scheid and Erilova, 2010). The production of seeds was highly variable in this study but was generally much greater than what has been reported previously in triploid cannabis (Table 1 and 2). Crawford *et al*. (2021) demonstrated that homogametic (XXX) triploid *C. sativa* pollinated by a homogametic (XX) diploid parent and homogametic (XXXX) tetraploid parent treated with STS produced 8.25 and 17.25 total seeds on average respectively. When the seed parent was a homogametic diploid (XX), the plants produced 354 and 232.75 seeds per plant representing a 97.7% and 32.3% reduction in seed production respectively (Crawford *et al*., 2021). Kurtz, Brand and Lubell-Brand (2024) also found a 99.5% reduction in seeds when triploids were pollinated by a diploid. Moreover, Suchoff *et al.,* (2024) further demonstrated that triploids produced between 87% and 77% less seeds when compared to their diploid counterparts. In this study, homogametic (XXX) triploids produced 147.3 ± 58.66 seeds on average (Table 2). However, the number of seeds per plant varied dramatically in this study, demonstrating that seedlessness is highly variable, even within a single population. This is also consistent with other triploid genotypes produced in our lab that have been nearly seedless as reported by Crawford *et al.,* (2021)(unpublished data). As such, it appears that fertility is highly variable in triploid *Cannabis* and likely depends upon the genetic background of the plants.

Little has been studied regarding the yield of male *C. sativa* plants. Moreover, triploid cannabis studies have been focused on triploid female (XXX) flower yields (Bagheri & Mansouri, 2015; Parsons *et al*., 2019; Crawford *et al*., 2021; Fernandes *et al*., 2023). In this study, it was observed that fresh weight increased linearly from male to female phenotypes (Table 1). Moreover, homogametic (XXX) plants significantly more fresh weight than heterogametic (XXY) (Table 2). However, this trend was not observed in dry total weight nor dried flower weight. This is likely due to the floral architecture when determining male and female cannabis plants. Male inflorescences are smaller relative to females and would ultimately translate to less dried weight (Raman *et al*., 2017). Moreover, due to the fragile nature of male cannabis inflorescences, many flowers were knocked off and not included in the final dry weight measurements.

Dioecious species such as *Juniperus communis* and *Podocarpus nagi* display enhanced male vigor (Nanami, Kawaguchi, and Yamakura, 2005; Zeidler *et al*., 2020). Male cannabis plants have also been reported to grow taller relative to female plants (Ramen *et al*., 2017). In this study, predominately female monoecious phenotypes produced similar fresh, and dry weights relative to female phenotypes (Table 1). However, it was observed that the male phenotypes produced a significantly lower HI relative to the predominantly female monoecious (Table 1). This suggests that phenotypic males produce greater amounts of vegetative biomass than reproductive relative to phenotypic females and monoecious types.

There have been minimal studies aimed at examining male *C. sativa* THC production. Male cannabis plants typically yield low amounts of THC and are usually discarded in commercial facilities producing for the recreational market (Raman *et al*., 2017; Small, 2017). However, Mansouri & Bagheri (2017) found that THC was reduced in tetraploid male flowers of hemp plants. In this study, phenotypic females had significantly higher THC relative to the males, but not predominant female monoecious phenotypes. This is likely due to the classification used of females, males, and monoecious plants. Phenotypic females exclusively produce bracts that are typically covered with glandular trichomes (Small, 2017). Phenotypic males produce inflorescences typically with few glandular trichomes. Thus, XXY individuals phenotypically classed as male were included in the comparison. The male floral organs found on the monoecious individuals likely reduced the amount of trichomes produced in the inflorescence thereby reducing the THC content. Similarly, the visual classification system of the phenotypes is based on the amount of male inflorescence. Since male flowers produce less trichomes than female, naturally as the phenotypes became more masculine, average THC and THCA decreases.

There have minimal studies examining the average seed weight of homogametic (XXX) *C. sativa*. Suchoff *et al.,* (2024) found that triploids produced lighter seeds than their diploid counterparts. The group found triploid seeds were on average 12.53 mg and 12.50 mg where diploids were 13.80 mg and 13.86 mg based on cultivar respectively (Suchoff *et al*., 2024). In addition, a 2021 study of 6 diploid monoecious hemp varieties found that average seed weight was between 6.17 mg to 8.99 mg depending on the year (Tsaliki *et al*., 2021). This is lower than the average seed weight in this studies population (6.08 mg), However, the average seed weight was variable and dependent on cultivar (Tsaliki *et al*., 2021). While drug-type *C. sativa* cultivars do not see comparable average seed weights at a higher ploidy level, hemp cultivars that have been bred for seed production may have different results. To the author’s knowledge there are also no studies examining the yield of heterogametic (XXY) seed yield in any species. This study found significant differences in the average seed weight between both genotypes, and two phenotypes (Table 1 and 2). The higher average seed weight of XXX and phenotypic female plants relative to the XXY and predominantly male monoecious phenotypic plants may be due to the lack of complete female floral organs. Many monoecious phenotypes produced incomplete or partial male and female flowers (Fig. 5). These abnormalities in the inflorescence may have prevented normal seed development.

## Conclusion

This study represents one of the building blocks in understanding the deeper mechanisms behind sex determination in *C. sativa* and provides insight into the floral reversion of cannabis and its sexual plasticity. The results of this study point away from a strict Y and X-to-autosome ratio system of sex determination in *C. sativa* and suggest it is more complex. The production of monoecious plants with varying degrees of female to male flowers indicates that sex expression in *C. sativa* is tightly linked to endogenous hormonal production controlled by genes not confined to the sex chromosomes. Moreover, these results display that some monoecious heterogametic phenotypes produce similar amounts of biomass, cannabinoids, and seed relative to homogametic plants. In addition, the results indicate that triploid *C. sativa* is genotype-specific and highly variable in terms of its ability to produce seeds. Ultimately, this information may aid in developing plants that are less prone to producing male flowers and reduce unwanted pollination events.

## Acknowledgements

We acknowledge the support of the Natural Sciences and Engineering Research Council of Canada (NSERC) and Dycar Pharmaceuticals.

## Author contributions

Conceptualization: N.P, B.Y, A.M, B.H, D.T, A.M.P.J

Data curation: N.P, B.Y

Formal analysis: N.P

Funding acquisition: A.M.P.J

Investigation: N.P, B.Y, B.H, M.B

Methodology: N.P, B.Y

Project administration: A.M.P.J

Resources: A.M.P.J

Software: N.P

Supervision: A.M.P.J

Validation: N.P, B.Y

Visualization: N.P

Writing – original draft: N.P

Writing – review & editing: N.P, B.Y, A.M, B.H, D.T, A.M.P.J

## Data availability

Datasets used in this study are available upon request. Code used to analyze data in R studio is available upon request. Requests can be made to.

## Reference

Acquaah, G. (2012) ‘Polyploidy in Plant Breeding’, in Principles of Plant Genetics and Breeding. Wiley, pp. 452–469. Available at: 10.1002/9781118313718.ch24.

Adal, A.M. et al. (2021) ‘Comparative RNA-Seq analysis reveals genes associated with masculinization in female Cannabis sativa’, Planta, 253(1), p. 17. Available at: 10.1007/s00425-020-03522-y.

Alonso, J.M. and Stepanova, A.N. (2004) ‘The Ethylene Signaling Pathway’, Science, 306(5701), pp. 1513–1515. Available at: 10.1126/science.1104812.

Anamthawat-Jónsson, K. et al. (2021) ‘Naturally occurring triploid birch hybrids from woodlands in Iceland are partially fertile’, New Forests, 52(4), pp. 659–678. Available at: 10.1007/s11056-020-09816-z.

Bagheri, M. and Mansouri, H. (2015) ‘Effect of Induced Polyploidy on Some Biochemical Parameters in Cannabis sativa L.’, Applied Biochemistry and Biotechnology, 175(5), pp. 2366–2375. Available at: 10.1007/s12010-014-1435-8.

Balant, M. et al. (2022) ‘Novel Insights into the Nature of Intraspecific Genome Size Diversity in Cannabis sativa L.’, Plants, 11(20). Available at: 10.3390/plants11202736.

Baldini, M. et al. (2018) ‘The Performance and Potentiality of Monoecious Hemp (Cannabis sativa L.) Cultivars as a Multipurpose Crop’, Agronomy, 8(9), p. 162. Available at: 10.3390/agronomy8090162.

Bhalerao, R.P. and Bennett, M.J. (2003) ‘The case for morphogens in plants’, Nature Cell Biology, 5(11), pp. 939–943. Available at: 10.1038/ncb1103-939.

Borin, M. et al. (2021) ‘Developing and Testing Molecular Markers in Cannabis sativa (Hemp) for Their Use in Variety and Dioecy Assessments’, Plants 2021, Vol. 10, Page 2174, 10(10), p. 2174. Available at: 10.3390/PLANTS10102174.

Boualem, A. et al. (2015) ‘A cucurbit androecy gene reveals how unisexual flowers develop and dioecy emerges’, Science, 350(6261), pp. 688–691. Available at: 10.1126/science.aac8370.

Clare, S.J. et al. (2024) ‘An affordable and convenient diagnostic marker to identify male and female hop plants’, G3: Genes, Genomes, Genetics, 14(1). Available at: 10.1093/g3journal/jkad216.

Crawford, S. et al. (2021) ‘Characteristics of the Diploid, Triploid, and Tetraploid Versions of a Cannabigerol-Dominant F1 Hybrid Industrial Hemp Cultivar, Cannabis sativa “Stem Cell CBG”’, Genes, 12(6), p. 923. Available at: 10.3390/genes12060923.

Crow, J.F. (1994) ‘Hitoshi Kihara, Japan’s pioneer geneticist.’, Genetics, 137(4), pp. 891–894. Available at: 10.1093/genetics/137.4.891.

Dellaporta, S.L. and Calderon-Urrea, A. (1993) ‘Sex determination in flowering plants.’, The Plant Cell, 5(10), pp. 1241–1251. Available at: 10.1105/tpc.5.10.1241.

Dubois, M., Van den Broeck, L. and Inzé, D. (2018) ‘The Pivotal Role of Ethylene in Plant Growth’, Trends in Plant Science, 23(4), pp. 311–323. Available at: 10.1016/j.tplants.2018.01.003.

Fernandes, H.P. et al. (2023) ‘Cultivar-dependent phenotypic and chemotypic responses of drug-type Cannabis sativa L. to polyploidization’, Frontiers in Plant Science, 14. Available at: 10.3389/fpls.2023.1233191.

Flajšman, M., Slapnik, M. and Murovec, J. (2021) ‘Production of Feminized Seeds of High CBD Cannabis sativa L. by Manipulation of Sex Expression and Its Application to Breeding’, Frontiers in Plant Science, 12(November), pp. 1–12. Available at: 10.3389/fpls.2021.718092.

Galoch, E. (1978) ‘The hormonal control of sex differentiation in dioecious plants of hemp (Cannabis sativa). The influence of plant growth regulators on sex expression in male and female plants’, Acta Societatis Botanicorum Poloniae, 47(1–2), pp. 153–162. Available at: 10.5586/asbp.1978.013.

Ghosh, D. et al. (2023) ‘Monoecious Cannabis sativa L. discloses the organ-specific variation in glandular trichomes, cannabinoids content and antioxidant potential’, Journal of Applied Research on Medicinal and Aromatic Plants, 35, p. 100476. Available at: 10.1016/j.jarmap.2023.100476.

Grant, S. et al. (1994) ‘Genetics of sex determination in flowering plants’, Developmental Genetics, 15(3), pp. 214–230. Available at: 10.1002/dvg.1020150304.

Haunold, A. (1971) ‘ Cytology, Sex Expression, and Growth of a Tetraploid ✕ Diploid Cross in Hop ( Humulus lupulus L.) 1 ’, Crop Science, 11(6), pp. 868–871. Available at: 10.2135/cropsci1971.0011183x001100060031x.

Heslop-Harrison, J. (1956) ‘Auxin and Sexuality in Cannabis sativa’, Physiologia Plantarum, 9(4), pp. 588–597. Available at: 10.1111/j.1399-3054.1956.tb07821.x.

Jones, M. and Monthony, A.S. (2022) ‘Cannabis Propagation’, in Handbook of Cannabis Production in Controlled Environments. CRC Press, pp. 91–121. Available at: 10.1201/9781003150442-4.

Köhler, C., Mittelsten Scheid, O. and Erilova, A. (2010) ‘The impact of the triploid block on the origin and evolution of polyploid plants’, Trends in Genetics, 26(3), pp. 142–148. Available at: 10.1016/j.tig.2009.12.006.

Kurtz, L.E., Brand, M.H. and Lubell-Brand, J.D. (2020) ‘Production of Tetraploid and Triploid Hemp’, HortScience, 55(10), pp. 1703–1707. Available at: 10.21273/HORTSCI15303-20.

Kurtz, L.E., Brand, M.H. and Lubell-Brand, J.D. (2024) ‘Cannabis Triploids Exhibit Reduced Fertility and Similar Growth and Flower Production Compared to Diploids’, Journal of the American Society for Horticultural Science, 149(2), pp. 75–85. Available at: 10.21273/JASHS05359-23.

Lubell, J.D. and Brand, M.H. (2018) ‘Foliar sprays of silver thiosulfate produce male flowers on female hemp plants’, HortTechnology, 28(6), pp. 743–747. Available at: 10.21273/HORTTECH04188-18.

Mansouri, H. and Bagheri, M. (2017) ‘Induction of Polyploidy and Its Effect on Cannabis sativa L.’, in Cannabis sativa L. - Botany and Biotechnology. Cham: Springer International Publishing, pp. 365–383. Available at: 10.1007/978-3-319-54564-6_17.

Martin, A. et al. (2009) ‘A transposon-induced epigenetic change leads to sex determination in melon’, Nature, 461(7267), pp. 1135–1138. Available at: 10.1038/nature08498.

Martínez, C. and Jamilena, M. (2021) ‘To be a male or a female flower, a question of ethylene in cucurbits’, Current Opinion in Plant Biology, 59, p. 101981. Available at: 10.1016/j.pbi.2020.101981.

Mckay, H.L. (2021) A Morphological and Molecular Investigation into the Role of Ethylene in Cannabis sativa Sex Determination.

McLeod, A. et al. (2023) ‘In Vivo and In Vitro Chromosome Doubling of “I3” Hemp’, HortScience, 58(9), pp. 1018–1022. Available at: 10.21273/HORTSCI17169-23.

Mejía Londoño, H.A., Barrera-Sánchez, C.F. and Córdoba Gaona, O. de J. (2023) ‘Sex reversal in female cannabis plants as a in response to male flowering promoters’, Revista Facultad Nacional de Agronomía Medellín, 76(3), pp. 10427–10435. Available at: 10.15446/rfnam.v76n3.102852.

Mohan Ram, H.Y. and Sett, R. (1982) ‘Reversal of Ethephon-Induced Feminization in Male Plants of Cannabis sativa by Ethylene Antagonists’, Zeitschrift für Pflanzenphysiologie, 107(1), pp. 85–89. Available at: 10.1016/S0044-328X(11)80012-7.

Moher, M., Jones, M. and Zheng, Y. (2021) ‘Photoperiodic response of in vitro cannabis sativa plants’, HortScience, 56(1), pp. 108–113. Available at: 10.21273/HORTSCI15452-20.

Moliterni, V.M.C. et al. (2004) ‘The sexual differentiation of Cannabis sativa L.: A morphological and molecular study’, Euphytica, 140(1–2), pp. 95–106. Available at: 10.1007/s10681-004-4758-7.

Monthony, A.S. et al. (2021) ‘Flower power: floral reversion as a viable alternative to nodal micropropagation in Cannabis sativa.’, In Vitro Cellular and Developmental Biology - Plant, 57(6), pp. 1018–1030. Available at: 10.1007/s11627-021-10181-5.

Monthony, Adrian S., de Ronne, M. and Torkamaneh, D. (2024) ‘Exploring ethylene-related genes in Cannabis sativa: implications for sexual plasticity’, Plant Reproduction [Preprint]. Available at: 10.1007/s00497-023-00492-5.

Monthony, Adrian S, de Ronne, M. and Torkamaneh, D. (2024) ‘Exploring ethylene-related genes in Cannabis sativa: implications for sexual plasticity’, Plant Reproduction [Preprint]. Available at: 10.1007/s00497-023-00492-5.

Moon, Y.-H. et al. (2020) ‘Effect of Timing of Ethephon Treatment on the Formation of Female Flowers and Seeds from Male Plant of Hemp (Cannabis sativa L.)’, The Plant Resources Society of Korea, 33(6), pp. 682–688. Available at: 10.7732/kjpr.2020.33.6.682.

Mudge, E.M., Murch, S.J. and Brown, P.N. (2017) ‘Leaner and greener analysis of cannabinoids’, Analytical and Bioanalytical Chemistry, 409(12), pp. 3153–3163. Available at: 10.1007/s00216-017-0256-3.

Oda, A. et al. (2022) ‘Temporary Reduction and Control of Female Flower Expression in Cucumber (Cucumis sativus L.) by Application of 1-Methylcyclopropene’, Horticulture Journal, 91(1), pp. 42–48. Available at: 10.2503/hortj.UTD-307.

Owen, L.C., Suchoff, D.H. and Chen, H. (2023) ‘A Novel Method for Stimulating Cannabis sativa L. Male Flowers from Female Plants’, Plants, 12(19), p. 3371. Available at: 10.3390/plants12193371.

Parsons, J.L. et al. (2019) ‘Polyploidization for the genetic improvement of cannabis sativa’, Frontiers in Plant Science, 10. Available at: 10.3389/fpls.2019.00476.

Petersson, S. V. et al. (2009) ‘An Auxin Gradient and Maximum in the Arabidopsis Root Apex Shown by High-Resolution Cell-Specific Analysis of IAA Distribution and Synthesis’, The Plant Cell, 21(6), pp. 1659–1668. Available at: 10.1105/tpc.109.066480.

Philbrook, R. et al. (2023) ‘Naturally Occurring Triploidy in Cannabis’, Plants, 12(23). Available at: 10.3390/plants12233927.

Prentout, D. et al. (2020) ‘An efficient RNA-seq-based segregation analysis identifies the sex chromosomes of Cannabis sativa’, Genome Research, 30(2), pp. 164–172. Available at: 10.1101/gr.251207.119.

Punja, Z.K. and Holmes, J.E. (2020) ‘Hermaphroditism in Marijuana (Cannabis sativa L.) Inflorescences – Impact on Floral Morphology, Seed Formation, Progeny Sex Ratios, and Genetic Variation’, Frontiers in Plant Science, 11. Available at: 10.3389/fpls.2020.00718.

Raman, V. et al. (2017) ‘Morpho-Anatomy of Marijuana (Cannabis sativa L.)’, in Cannabis sativa L. - Botany and Biotechnology. Cham: Springer International Publishing, pp. 123–136. Available at: 10.1007/978-3-319-54564-6_5.

Sattler, M.C., Carvalho, C.R. and Clarindo, W.R. (2016) ‘The polyploidy and its key role in plant breeding’, Planta. Springer Verlag, pp. 281–296. Available at: 10.1007/s00425-015-2450-x.

Shephard, H.L. et al. (2000) ‘Sexual development and sex chromosomes in hop’, New Phytologist, 148(3), pp. 397–411. Available at: 10.1046/j.1469-8137.2000.00771.x.

Shi, J., Schilling, S. and Melzer, R. (2024) ‘Morphological and genetic analysis of inflorescence and flower development in hemp (Cannabis sativa L.)’. Available at: 10.1101/2024.01.25.577276.

Small, E. (2017) ‘Classification of Cannabis sativa L.: In relation to agricultural, biotechnological, medical and recreational utilization’, in Cannabis sativa L. - Botany and Biotechnology. Springer International Publishing, pp. 1–62. Available at: 10.1007/978-3-319-54564-6_1.

Small, E. (2022) ‘Genetics and Plant Breeding of Cannabis sativa for Controlled Environment Production’, in Handbook of Cannabis Production in Controlled Environments. CRC Press, pp. 41–90. Available at: 10.1201/9781003150442-3.

Small, E. and Jomphe, M. (1989) ‘A synopsis of the genus Medicago (Leguminosae)’, Canadian Journal of Botany, 67(11), pp. 3260–3294. Available at: 10.1139/b89-405.

Spitzer-Rimon, B. et al. (2019) ‘Architecture and florogenesis in female Cannabis sativa plants’, Frontiers in Plant Science, 10. Available at: 10.3389/fpls.2019.00350.

Subasinghe Arachchige, E.C.W., et al. (2022) ‘Morphological characteristics of pollen from triploid watermelon and its fate on stigmas in a hybrid crop production system’, Scientific Reports, 12(1), p. 3222. Available at: 10.1038/s41598-022-06297-2.

Suchoff, D.H. et al. (2024) ‘Characterization of agronomic performance and sterility in triploid and diploid cannabinoid hemp’, Agronomy Journal [Preprint]. Available at: 10.1002/agj2.21618.

Tang, Q. et al. (2023) ‘Transcriptomic and metabolomic analyses reveal the differential accumulation of phenylpropanoids and terpenoids in hemp autotetraploid and its diploid progenitor’, BMC Plant Biology, 23(1). Available at: 10.1186/s12870-023-04630-z.

Tolety, J. and Sane, A. (2011) ‘Antirrhinum’, in Wild Crop Relatives: Genomic and Breeding Resources. Berlin, Heidelberg: Springer Berlin Heidelberg, pp. 1–14. Available at: 10.1007/978-3-642-21201-7_1.

Törjék, O. et al. (2002) ‘Novel male-specific molecular markers (MADC5, MADC6) in hemp’, Euphytica, 127(2), pp. 209–218. Available at: 10.1023/A:1020204729122.

Tsaliki, E. et al. (2021) ‘Fibre and seed productivity of industrial hemp (Cannabis sativa l.) varieties under mediterranean conditions’, Agronomy, 11(1). Available at: 10.3390/agronomy11010171.

Zhang, X.-L. et al. (2016) ‘Comparative Proteomic Analysis of Mature Pollen in Triploid and Diploid Populus deltoides’, International Journal of Molecular Sciences, 17(9), p. 1475. Available at: 10.3390/ijms17091475.

